# Modelling the impact of tuberculosis preventive therapy: the importance of disease progression assumptions

**DOI:** 10.1101/666669

**Authors:** Tom Sumner, Richard G. White

**Affiliations:** TB Modelling Group, TB Centre, Centre for the Mathematical Modelling of Infectious Disease, Department of Infectious Disease Epidemiology, London School of Hygiene & Tropical Medicine, United Kingdom

## Abstract

**Background:** Following infection with Mycobacterium tuberculosis (*M.tb*) individuals may rapidly develop tuberculosis (TB) disease or enter “latent” infection state with a low risk of progression to disease. The mechanisms underlying this process are incompletely known. Mathematical models use a variety of structures and parameterisations to represent this progression from infection with *M.tb* to disease. This structural and parametric uncertainty may affect the predicted impact of interventions leading to incorrect conclusions and decision making.

**Methods:** We used a simple dynamic transmission model to explore the effect of uncertainty in model structure and parameterisation on the predicted impact of scaling up preventive therapy. We compared three commonly used model structures and used parameter values from two different data sources. Models 1 and 2 are equally consistent with observations of the time from infection to disease. Model 3, produces a worse fit to the data, but is widely used in published modelling studies. We simulated treatment of 5% of all *M.tb* infected individuals per year in a population of 10,000 and calculated the reduction in TB incidence and number needed to treat to avert one TB case over 10 years.

**Results:** The predicted impact of the preventive therapy intervention depended on both the model structure and the parameterisation of that structure. For example, at a baseline annual TB incidence of 500/100,000, the impact ranged from 11% to 27% and the number needed to treat to avert one TB case varied between 38 and 124. The relative importance of structure and parameters varied depending on the baseline incidence of TB.

**Discussion:** Our analysis shows that the choice of model structure and the parameterisation can influence the predicted impact of interventions. Modelling studies should consider incorporating structural uncertainty in their analysis. Not doing so may lead to incorrect conclusions on the impact of interventions.

## Introduction

Tuberculosis (TB) is an infectious disease most commonly caused by the bacteria mycobacterium tuberculosis (*M.tb*). Following infection with *M.tb* it is commonly stated that 10-15% of individuals will develop disease in their lifetime [1]. The risk of developing TB is known to vary with time since infection, with the highest risk in the first year following infection [2]. Disease which occurs soon after infection is typically referred to as primary TB. TB may also occur many years after initial exposure, either through reactivation of “latent” infection or due to re-infection [3, 4]. The mechanisms that underlie this process are incompletely known.

Mathematical models are increasingly used to predict the impact of TB control strategies and to inform policy making. These models use a variety of structures and parameters to represent the progression from infection with *M.tb* to TB disease. While modelling studies often explore the sensitivity of results to parameters, sensitivity to the choice of model structure is rarely addressed in a systematic way.

Two recent papers [5, 6] have compared different model structures to describe the progression from infection to disease and assessed their ability to reproduce observations of the time from *M.tb* infection to TB disease onset. These analyses suggested that certain model structures capture these dynamics better than others, and that the parameters and structures used in many published modelling studies do not accurately reproduce the temporal pattern of disease following exposure to *M.tb* observed in many settings. These findings raise important questions about how the structure and parameters commonly used to represent progression to disease may affect model predictions of the impact of control strategies.

In this paper we use a simple dynamic transmission model of TB preventive therapy to explore the effect of the model structure and parameterisation used to represent progression from infection to disease on the predicted impact of a control strategy. The World Health Organisation (WHO) recommends TB preventive therapy for a number of high risk populations including people living with HIV, household contacts of pulmonary TB cases and dialysis and organ transplant patients [7]. Previous modelling [8–10] has highlighted increased use of preventive therapy as a key component of reaching the WHO global TB targets [11]. As preventive therapy aims to prevent progression from infection to disease, it is important to understand how the model structure and parameters used to represent this process effect the model results.

## Methods

### Selection of model structures

We compare 3 model structures for progression from infection to disease that are commonly used in TB modelling studies. The systematic literature review reported in Menzies et al [6] found that these 3 structures accounted for approximately 70% of all published TB models. Both Menzies et al [6] and Ragonnet et al [5] showed that models 1 and 2 gave equally good fit to the available data on the cumulative incidence of TB following infection and it was not possible to distinguish a “best” model. Both studies also found model 3 gave a significantly worse fit, but as this structure is employed in a large proportion of published modelling studies (approximately 50% based on the literature review in [6]) we included it in our analysis.

### Description of models

Figure 1 shows the different model structures incorporated into a simple dynamic transmission model of TB. The equations and steady state solutions for each of the models are given in the appendix.

**Figure 1.**
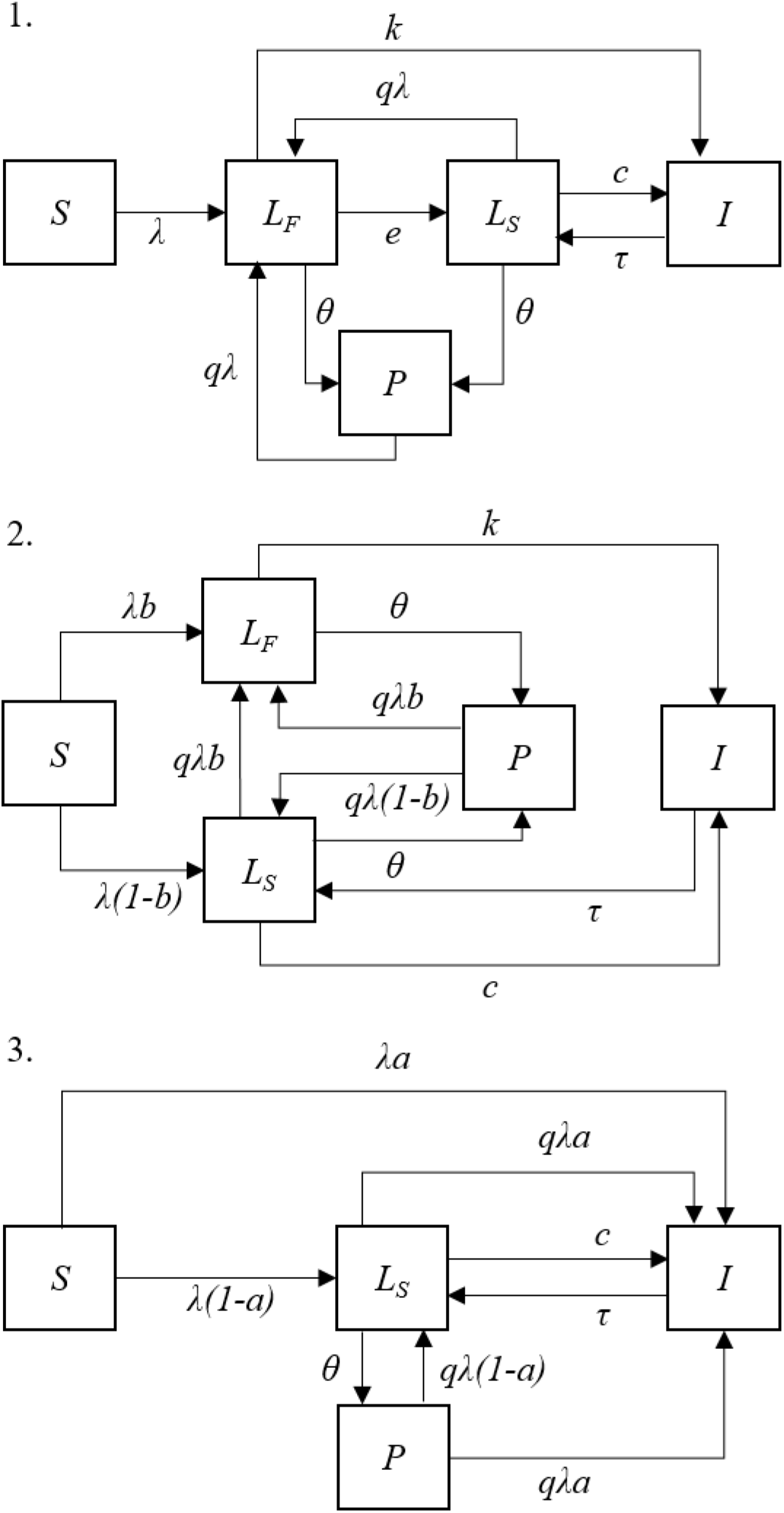
Schematic of model structures. *S* = susceptible; *L*_*F*_ = “fast” latent state; *L*_*S*_ = “slow” latent state; *I* = TB disease; *P* = post preventive therapy. Definitions of model parameters are given in table 1.

The following features are common to all models. Susceptible individuals (*S*) are infected with *M.tb* at a rate *λ*=*βI* where *β* is the rate of effective contact and *I* is the prevalence of TB. Background mortality is modelled at a constant rate, *u*, in all states. In addition, those in the disease state, *I*, are subject to an additional disease associated mortality rate, *m*. The birth rate is set to maintain a constant population size. All births are assumed to be susceptible. In all models we assume that those with TB disease (*I*) are removed back to the “slow” latent state (*L*_*s*_) at a rate, *τ*. This represents effective treatment and natural recovery from disease. Prior exposure to *M.tb*. is assumed to confer some immunity against re-infection. This is captured through the parameter *q* which represents the relative susceptibility to re-infection among those with latent infection compared to the susceptibility to first infection among previously uninfected individuals.

Preventive therapy is incorporated in a simple manner, aiming to be consistent across all models. The intention is not to make detailed predictions of the impact of preventive therapy, rather to illustrate how the impact may vary due to the choice of model structure. In each case we assume the population in all latent states is provided with effective preventive therapy at a constant rate *θ*. Those given preventive therapy move to the preventive therapy state, *P* where they have no risk of progression to disease but can be re-infected.

Model 1 consists of 2 sequential latent states. Following infection all individuals enter the “fast” latent state (*L*_*F*_) where they have an annual rate of progression to disease, *k*. Those who do not develop disease transition to the “slow” latent state (*L*_*S*_), at an annual rate, *e*, where they have an annual rate of disease progression *c* (where *c*<*k*). Biologically, this assumes that all infected individuals have the same risk of developing TB following infection. Individuals in the “slow” latent state (*L*_*s*_) and those who have been given preventive therapy (*P*) can be re-infected and return to *L*_*F*_.

Model 2 consists of 2 parallel latent states. Following infection, some proportion (*b*) enter the “fast” latent state (*L*_*F*_) where they have an annual rate of progression to disease, *k*. The remainder (1-*b*) enter the “slow” latent state (*L*_*S*_) where they have an annual rate of disease progression *c* (where *c*<*k*). Biologically, this assumes that some proportion of individuals (*b*) are pre-determined to have a high risk of developing TB following infection. Individuals in *L*_*S*_ and *P* can be re-infected with a proportion *b* moving to *L*_*F*_ and the remainder remaining in/going to *L*_*S*_.

Model 3 consists of a single “slow” latent state, *L*_*S*_. Following infection, some proportion (*a*) develop disease immediately. The remainder (1-*a*) enter the “slow” latent state where they have an annual rate of disease progression *c*. This can be seen as equivalent to model 2 but with an infinite rate of progression from the fast latent state to disease. Individuals in *L*_*S*_ and *P* can be re-infected with a proportion *a* progressing directly to disease and the remainder remaining in/going to *L*_*S*_.

### Model parameterisation

The analysis in Ragonnet et al [5] and Menzies et al [6] compared the various model structures to different data sets and therefore identified different best-fitting parameters. We compare the impact of using these 2 different parameter sources on the model predictions of the impact of preventive therapy. The parameter values are shown in table 1.

**Table 1.** Parameters for the transmission model. ^$^For Ragonnet et al we use the parameters estimated using least squares methods. *These parameters are not reported in Ragonnet et al so have been estimated to reproduce figure S14. Parameters in Ragonnet et al are reported in daily units and have been converted to annual units.

|  | Model |  |  |  |  |  |
| --- | --- | --- | --- | --- | --- | --- |
|  | 1 |  | 2 |  | 3 |  |
|  | Menzies | Ragonnet<sup>§</sup> | Menzies | Ragonnet<sup>§</sup> | Menzies | Ragonnet<sup>§</sup> |
| $a$, proportion progressing directly to disease | - | - | - | - | 0.0665 | 0.085* |
| $b$, proportion entering fast latent state | - | - | 0.0866 | 0.09 | - | - |
| $c$, rate of progression to disease from slow latent state (per year) | $5.94 \times 10^{-4}$ | $2.01 \times 10^{-3}$ | $5.94 \times 10^{-4}$ | $2.01 \times 10^{-3}$ | $3.37 \times 10^{-3}$ | $3.29 \times 10^{-3}$ |
| $k$, rate of progression to disease from fast latent state (per year) | 0.0826 | 0.4015 | 0.955 | 4.015 | - | - |
| $e$, rate of movement from fast latent state to slow latent state (per year) | 0.872 | 4.015 | - | - | - | - |
| $q$, relative susceptibility to re-infection compared to first infection | 0.5 | | | | | |
| $\tau$, rate of recovery from TB disease (per year) | 1 | | | | | |
| $\beta$, number of effective contacts (per year) | Varied to produce different TB incidence | | | | | |
| $m$, rate of TB associated mortality | 0.03 | | | | | |

### Modelling the impact of preventive therapy

To explore the effect of structure and parameterisation on the model predictions we simulated the introduction of preventive therapy for all latently infected individuals in a population with TB prevalence in steady state (see appendix for steady state solutions for each model). We explore a range of baseline TB incidence from 0 to 1000/100,000. We assumed an annual coverage of 100% efficacious preventive therapy of 5%. We calculated the percentage reduction in TB incidence (compared to the endemic equilibrium) after 10 years of preventive therapy. We also calculated the cumulative number of cases averted, the cumulative number of people given preventive therapy and the average number needed to treat (NNT) with preventive therapy to avert one TB case over the 10- year period assuming a constant population of 10,000.

## Results

### Cumulative risk of TB

Figure 2 and table 2 show the predicted cumulative incidence of TB after infection (in the absence of reinfection or preventive therapy) predicted by each model using each parameter set in table 1. Models 1 and 2 predict the same cumulative incidence over time. Beyond the time period of the data the models were fitted to (5 years and 10 years respectively, shown by the vertical dashed lines in figure 2), model 3 predicts a higher cumulative risk of TB than models 1 and 2.

**Table 2.** Cumulative proportion developing TB for each model and each parameter source.

|  | Menzies | Ragonnet |
| --- | --- | --- |
|  | Cumulative proportion of those infected developing TB |  |
| **Model 1** | 0.11 | 0.17 |
| **Model 2** | 0.11 | 0.17 |
| **Model 3** | 0.20 | 0.21 |
|  | Proportion of disease occur due to fast progression |  |
| **Model 1** | 0.76 | 0.52 |
| **Model 2** | 0.76 | 0.52 |
| **Model 3** | 0.33 | 0.40 |

**Figure 2.**
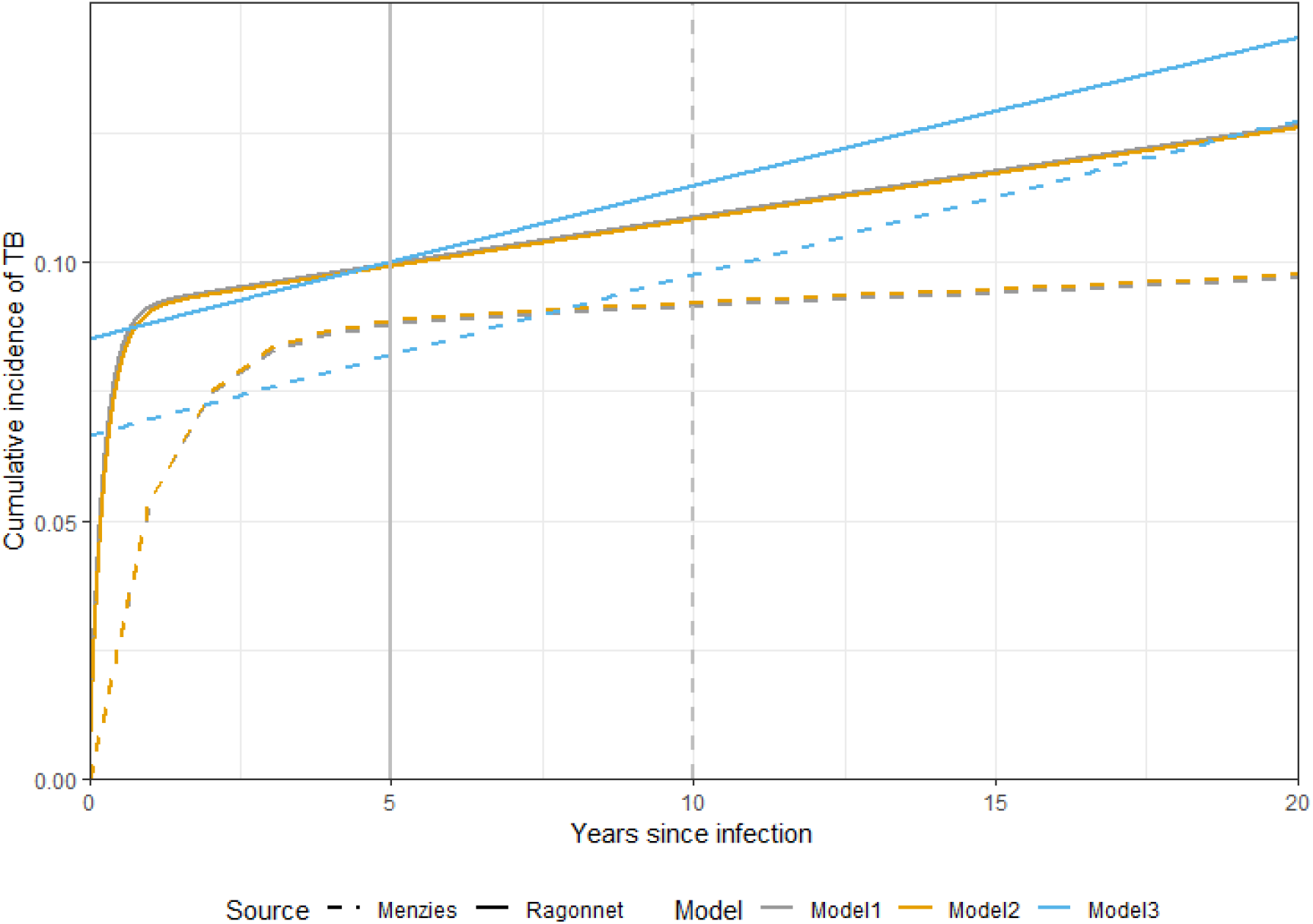
Cumulative proportion that have developed TB by time since infection. Colours indicate model, line type indicates source of parameter estimates. The vertical lines indicate the maximum time of the data to which the models were fitted. Note, results for models 1 and 2 overlap.

To better explore the relative effect of model structure and parameters we re-parameterised model 3 to give the same cumulative incidence as models 1 and 2. We fixed the rate of progression to disease from the slow latent state (*c*) to be the same in model 3 as in models 1 and 2 and calculated the value of *a*, the proportion progressing directly to disease in model 3, required to give the same cumulative incidence (see appendix equation 39). This gives new values for *a* of 0.084 for Menzies and 0.090 for Ragonnet. In the following analysis we consider both the original and updated parameterisations of model 3.

### Steady state incidence

Figure 3 shows the steady state TB incidence for each model as a function of the contact parameter, *β*. The left hand panel shows the results using the parameter estimates from Menzies et al, the right hand panel those obtained with the parameters from Ragonnet et al.

**Figure 3.**
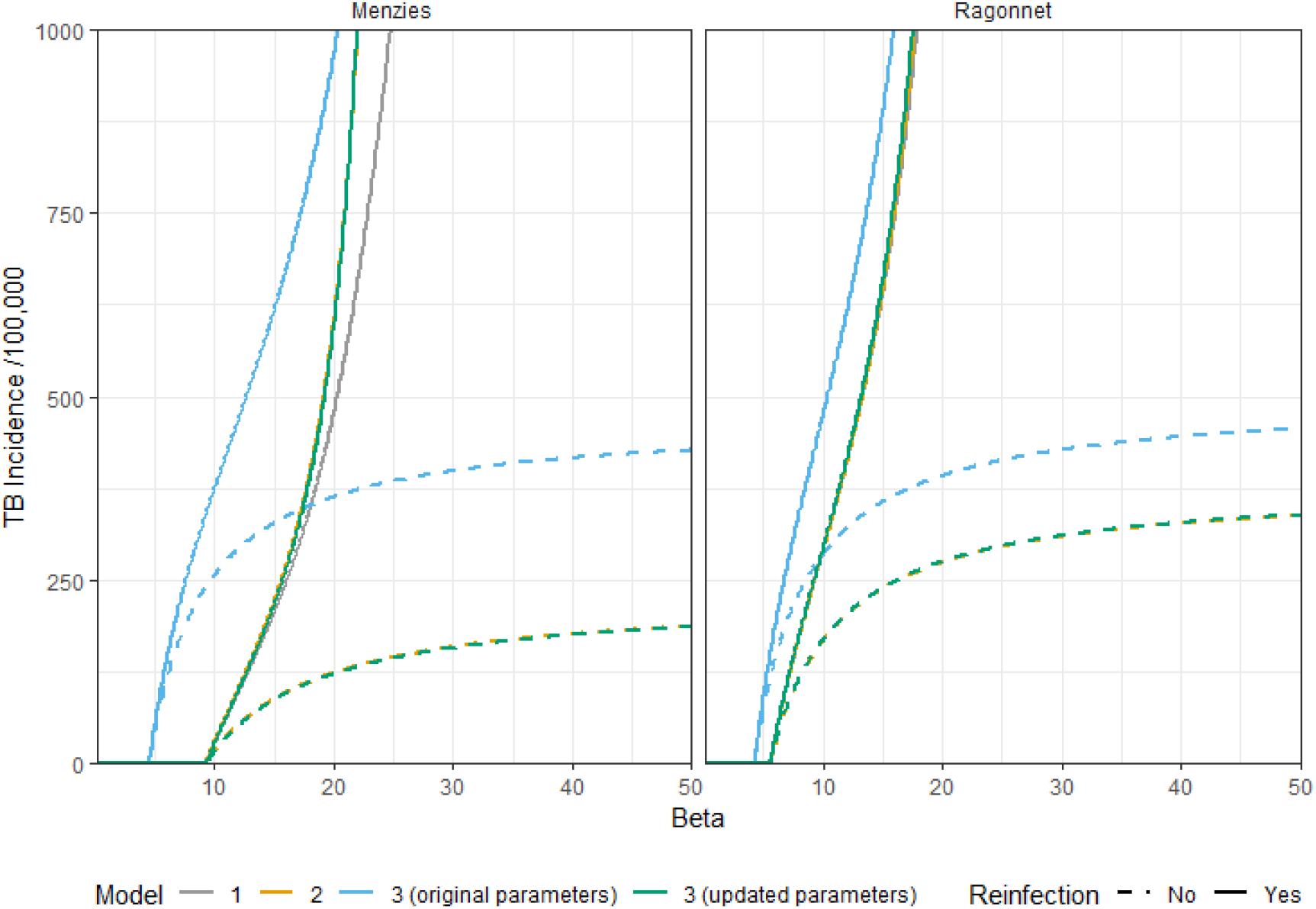
Steady state TB incidence as a function of the contact parameter, *β*. Left: results using parameter estimates from Menzies et al; right: results using estimates from Ragonnet et al. Colours indicate the different models. Dashed lines show the results when reinfection is not included in the model. Note, results for models 1, 2 and 3 (updated parameters) overlap.

In the full model including reinfection (solid lines in figure 3) the original parameterisation of model 3 (blue line) results in the highest incidence at a given value of *β* due to the higher lifetime risk of developing TB following infection (see figure 2). At low values of *β* models 1, 2 and the updated parameterisation of model 3 give the same incidence. However, as the contact rate, *β*, increases the model predictions diverge with a lower incidence predicted with model 1 at a given value of *β*. This divergence occurs due to differences in the risk of reinfection. In model 1, all individuals spend some time in the “fast” latent state where they are not at risk of reinfection and therefore the incidence of disease is lower. This role of reinfection can be seen by contrasting with the results from a model with no reinfection (dashed lines in figure 3). In this case models 1, 2 and the updated parameterisation of model 3 predict the same incidence as a function of *β*.

The same qualitative patterns are observed for both parameter sources but the divergence at higher incidence is smaller when using the parameters from Ragonnet et al.

### Impact of preventive therapy

Figure 4 shows the results of simulating 10 years of preventive therapy for each model structure as a function of the steady state TB incidence. Solid lines show the results using the parameterisation from Ragonnet et al., dashed lines the results using parameters from Menzies et al.

**Figure 4.**
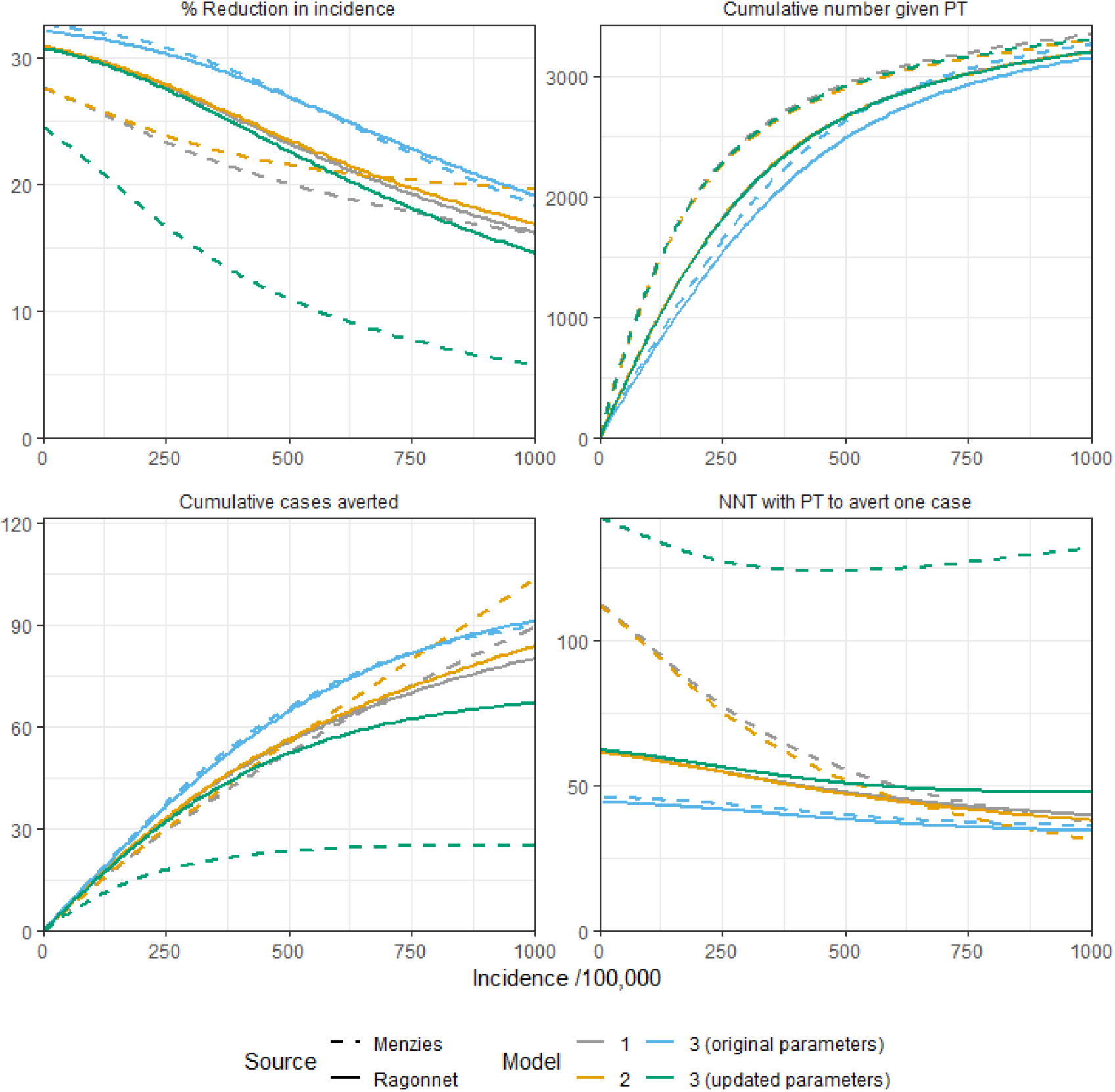
Results of simulating 10 years of preventive therapy as a function of steady state TB incidence. Top left: Percentage reduction in TB incidence from steady state equilibrium; top right: cumulative number given preventive therapy (assuming a constant population size of 10,000); bottom left: cumulative number of TB cases averted; bottom right: average number needed to treat with preventive therapy to avert one case of TB. Colours indicate the different models. Line types indicate the different sources of parameter estimates.

There is considerable difference between the predictions using different models and parameterisations. For example, at an incidence of 500/100,000 the predicted reduction in incidence (top left panel of figure 4) ranges from 11% to 27%. At the same incidence, the NNT (bottom left panel of figure 4) varies from 38 to 124.

For all models the impact declines as a function of increasing steady state TB incidence. This trend is due to two factors: at higher incidence the prevalence of infection is higher so there is less indirect benefit of reducing future transmission by preventing incident TB; at higher incidence the risk of reinfection after preventive therapy is greater which reduces the long-term benefit of treatment.

The largest impact is predicted using the original parameterisation of model 3. This is due to the higher risk of developing disease following infection in this model (see figure 2). As a result, the contact rate needed to produce a given incidence is smaller (see figure 3) and therefore the risk of reinfection after preventive therapy is lower. The prevalence of infection at a given incidence is also lower in this model which results in a greater indirect benefit of preventing incident TB. In contrast, the lowest impact is observed with the re-parameterised version of model 3. This is because it is not possible to directly prevent fast progression to disease in this model structure by providing preventive therapy to the latent populations; these cases do not pass through a “fast” latent state where they can be treated with preventive therapy. At low incidence we see a very similar predicted impact for models 1 and 2, however the predictions diverge at higher incidence, with a larger impact observed with model 2 compared to model 1. This divergence is due to the different risk of reinfection after preventive therapy (with no reinfection models 1 and 2 predict the same impact at any incidence). In model 1 a higher *β* value is needed to give the same incidence of disease (see figure 3) and therefore the risk of reinfection is higher at a given incidence. These qualitative patterns are the same for both parameter sources.

Despite the declining impact, the absolute number of cases averted increases with increasing steady state incidence because there are more cases which can be prevented. Similarly, the number of people treated with preventive therapy also increases with steady state incidence, reflecting the higher prevalence of latent infection. For models 1 and 2 the NNT declines with increasing steady state incidence while for model 3 we observe non-monotonic behaviour where the NNT initially declines with increasing incidence before increasing at higher incidence. This occurs because at a higher incidence a larger proportion of disease is due to reinfection and rapid progression. As this cannot be prevented in model 3 the impact predicted by model 3 declines more rapidly with increasing incidence. This behaviour was observed in a previous analysis using model with structure 1 [12]. We explore this further below.

### Comparison of models 1 and 2

The previous analysis of model structures for progression from infection to disease [5, 6] showed that it was not possible to select a “best” model using available data on timing of disease after infection and that models 1 and 2 produced an equally good fit. Figure 5 shows the results of combining the predictions of these 2 models only (i.e. excluding model 3 which gives a poorer fit to the data) within and across parameter sources. The solid black lines show the total envelope of predicted results while the shaded areas show the prediction when the results for both structures are combined for a given parameters source. Excluding model 3 reduces the overall range in model predictions however there is still considerable variation, particularly in the estimated NNT at low to intermediate incidence (right panel of figure 5). Figure 5 also highlights that the relative importance of model structure and parameters depends on the baseline incidence. At low to intermediate incidence the model structure does not affect the results and the uncertainty is almost entirely due to the different parameter sources. However, at higher incidence, the model structure becomes increasingly important.

**Figure 5.**
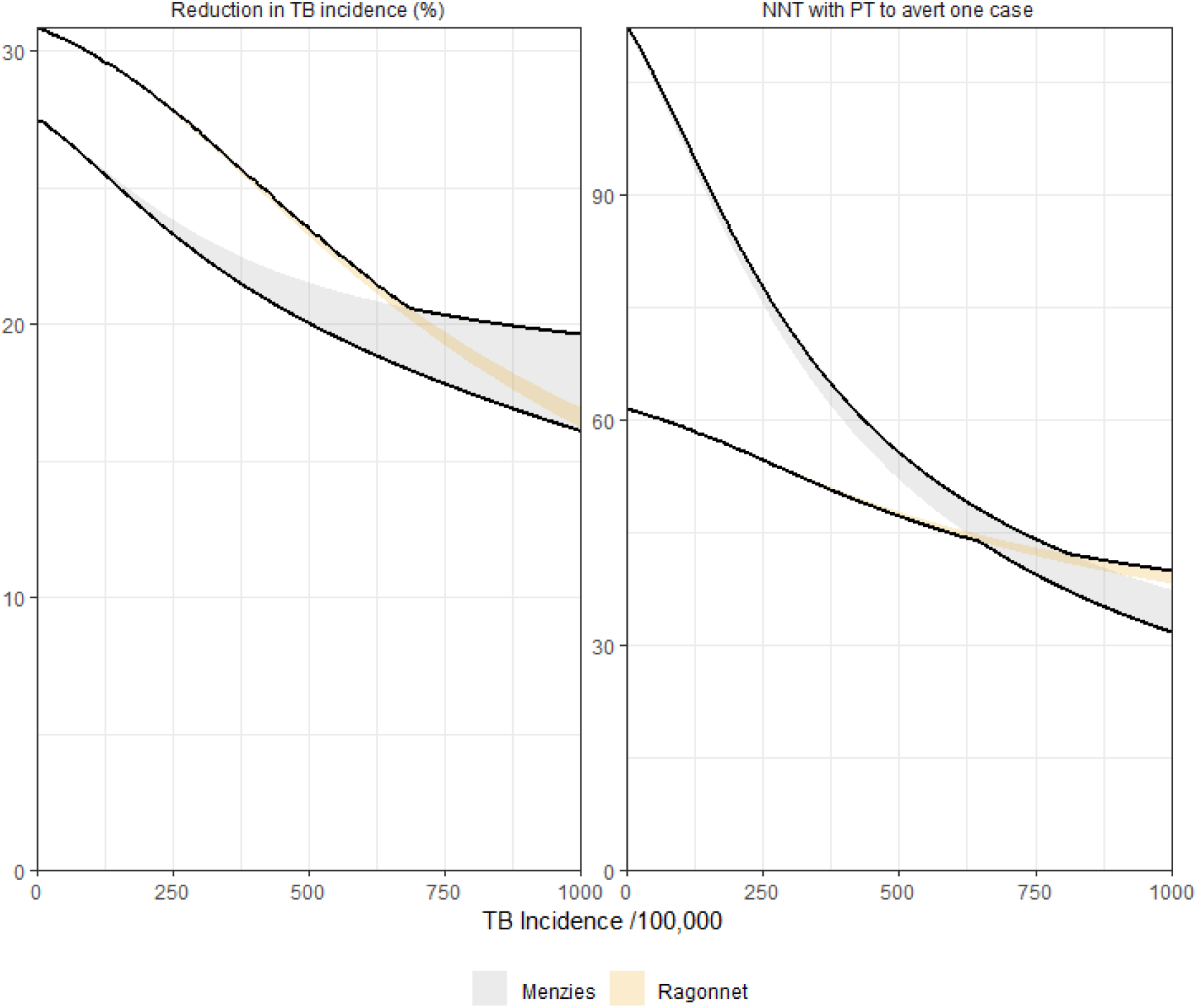
Range of predictions from models 1 and 2. Solid black lines show the total envelope of predictions for both models and both parameterisations. The shaded grey area shows the range using parameters from Menzies et al, the shaded yellow area the values using parameters from Ragonnet et al.

### Non-monotonic relationship between NNT and steady state incidence

Previous analysis of a simple model of preventive therapy [12] found a non-monotonic relationship between baseline incidence and NNT using model structure 1 with a minimum NNT occurring at an incidence in the region of 700/100,000. In this analysis we observed this non-monotonic relationship for model 3 but not for models 1 or 2 within the range of incidence explored. The key difference between the parameterisations of model 1 here and the analysis in [12] is the duration of the fast latent state which is determined by *e*, the rate of movement from the fast latent state to the slow latent state. In [12] this was assumed to be 5 years while in the parameterisations used here it is much shorter (between 0.25 and 1.15 years).

To explore this, we compared model 1 with values of *e* of 1, 0.5, 0.3̇33, 0.25 and 0.2 which give durations of the fast latent state of 1 to 5 years. We set the rate of progression from the slow latent state *c* = 5.94×10^−4^ and calculated the value of *k* (the rate of progression from the fast latent state) to give a cumulative risk of TB of 0.11, the same as with the parameters estimated in Menzies et al. (see table 2).

Figure 6 shows the predicted outputs of the simulated preventive therapy intervention for each duration of the “fast” latent state.

**Figure 6.**
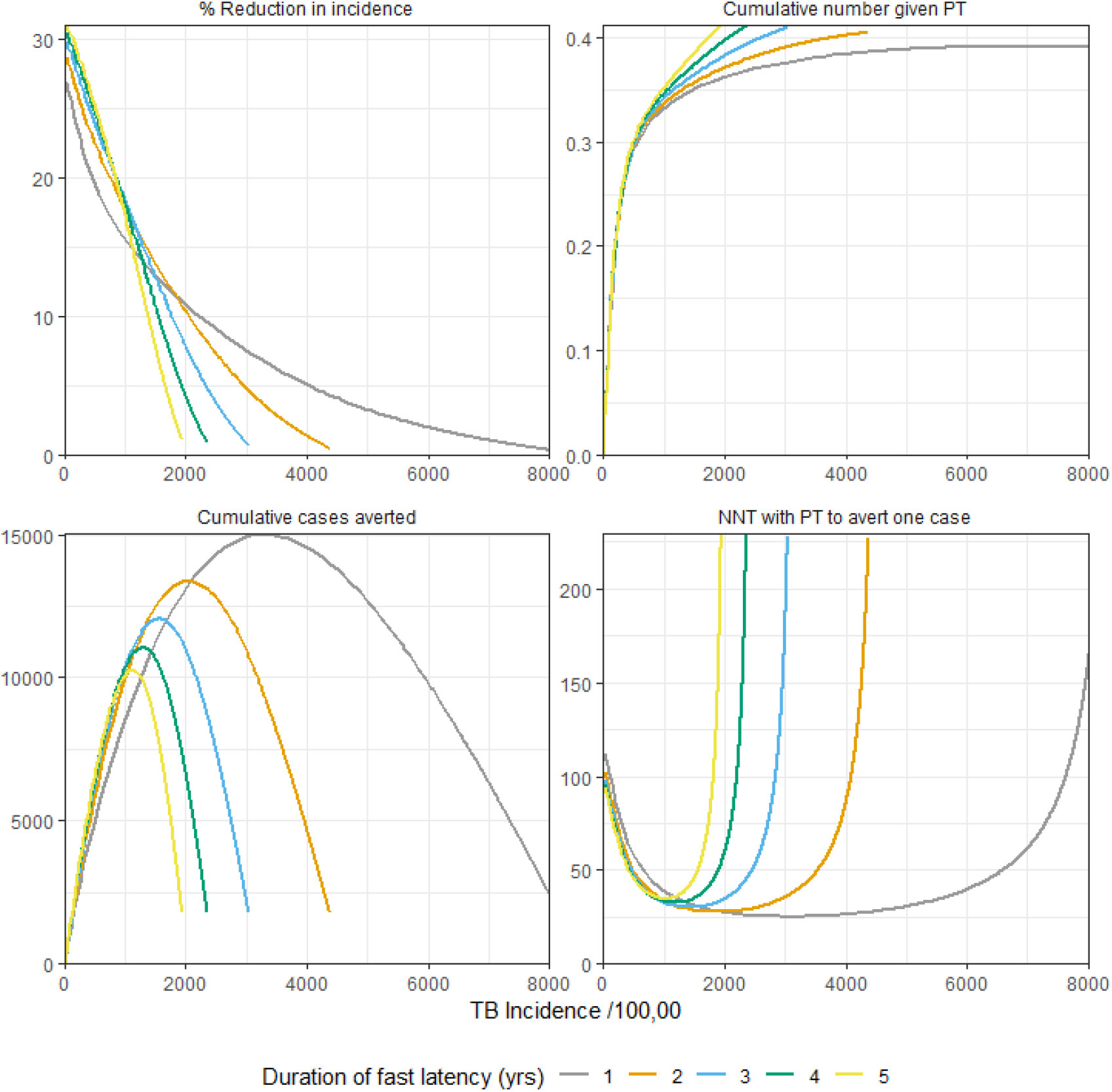
Predicted impact of preventive therapy in model 1 with different durations of “fast” latency. Top left: Percentage reduction in TB incidence from steady state equilibrium; top right: cumulative number given preventive therapy (assuming a constant population size of 10,000); bottom left: cumulative number of TB cases averted; bottom right: average number needed to treat with preventive therapy to avert one case of TB. Colours indicate the different durations of the “fast” latent state.

This shows that the non-monotonic behaviour is observed but that the “optimum” incidence (i.e. the value at which the NNT is minimised) is dependent on the duration of the fast latent state. With a duration of 5 years (yellow line) the minimum NNT occurs at an incidence of 1008/100,000. However, with a duration of 1 year (grey line) the minimum NNT occurs at a much higher incidence of 3685/100,000.

## Discussion

Our results show that both the model structure and parameter values used to represent progression from infection to disease can affect model predictions of the impact of preventive therapy on TB incidence. This highlights the importance of including both parametric and structural uncertainty in TB modelling studies. Failure to do so may result in inaccurate predictions of the potential impact of interventions.

Our analysis extends the findings of [5, 6] and shows that, in addition to producing a worse fit to data on the cumulative incidence of TB following infection, model 3 also predicts very different effects of preventive therapy when compared to models 1 and 2. Depending on the parameterisation of this model, it can both over or under estimate the impact compared to models 1 and 2. Our findings also show that models 1 and 2 (which fit the data equally well) can give different predictions of intervention impact.

We found that the relative importance of the model structure and the choice of parameters depends on the baseline incidence with the choice of structure becoming more important at higher incidence. This again highlights the need to consider both forms of uncertainty in modelling studies.

This work also suggests that the “optimum” incidence for use of preventive therapy found in [12] depends strongly on the model structure and parameterisation. Using the same model structure as in [12] we found that the NNT does not display non-monotonic behaviour within a plausible range of incidence unless the duration of the fast latent state is assumed to be much longer than estimated in [5, 6].

To allow us to explore a number of different model structures for the progression from infection to latency the rest of the model was kept as simple as possible. These simplifications may affect our findings. We used a very simple representation of demography, assuming a constant population size and a constant life expectancy. Previous work [13] has shown the importance of considering realistic age structure in models of TB transmission.

We assumed TB was in equilibrium before the introduction of preventive therapy, but trends in disease may affect the prevalence of infection and the contribution of ongoing transmission and reactivation to TB incidence. These factors are likely to influence the model predictions of intervention impact and may also affect the interaction between structure, parameters and the model outputs.

The representation of the preventive therapy intervention was greatly simplified to explore the impact of model structure on the results and the findings should not be interpreted as predictions of the likely impact of preventive therapy. As such we did not consider uncertainty in other parameters or assumptions describing the intervention. These are also likely to result in uncertainty in model predictions.

This work has focussed on the effect of progression model structure and parameters on the impact of preventive therapy. These features of TB models may also affect the predicted impact of other interventions. They may be particularly relevant in models used to study the impact of other preventive strategies such as vaccines but are likely to be important when considering a wide range of interventions.

## Conclusion

Uncertainty in model structure is often ignored in TB modelling studies. Future studies should aim to compare different structures when predicting impact of interventions to ensure the uncertainty in model predictions is captured more accurately.

To ensure both structural and parametric uncertainty can be explored in a systematic way, standardised methods for incorporating model structure are needed. Approaches such as Bayesian model averaging may be of value, but more work is required to increase their use in infectious disease modelling.

In addition to incorporating uncertainty in model predictions it is also important to understand how different sources of input uncertainty contribute to the variability in outputs. Methods to conduct quantitative sensitivity analysis of both model structure and parameters are needed. These would allow the key drivers of uncertainty to be identified and focus efforts on collecting data which will reduce uncertainty in future model predictions.

## Author Contributions

TS and RW conceived the study. TS conducted the analysis and drafted the manuscript. All authors contributed to revision of the manuscript and approved the final version.

## Appendix

The appendix provides further details of the model equations and steady state solutions.

### Model Equations

**Table A1.** Model variables.

| Variable | Description |
| --- | --- |
| $S$ | Susceptible. Never exposed to *M.tb.* |
| $L_F$ | Latently infected with *M.tb.* At risk of “fast” progression to active TB |
| $L_S$ | Latently infected with *M.tb.* At risk of “slow” progression to active TB |
| $I$ | Active TB disease |
| $P$ | Post-preventive therapy |
| $T$ | Total population |

**Table A2.** Model parameters.

| Parameter | Description |
| --- | --- |
| $a$ | Proportion progressing directly to disease after infection (model 3) |
| $b$ | Proportion entering fast latent state after infection (model 2) |
| $c$ | Rate of progression to disease from slow latent state (per year) (all models) |
| $e$ | Rate of movement from fast latent state to slow latent state (per year) (model 1) |
| $k$ | Rate of progression to disease from fast latent state (per year) (models 1 and 2) |
| $m$ | Excess mortality rate due to active TB (per year) |
| $q$ | Relative susceptibility to reinfection if previously infected |
| $u$ | Background mortality rate (per year) |
| $\lambda = \beta I$ | Force of infection |
| $\beta$ | Effective contact rate (per year) |
| $\theta$ | Rate of starting preventive therapy (per year) |

#### Model 1

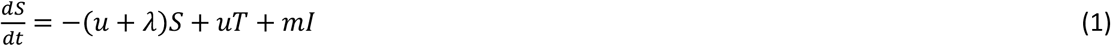

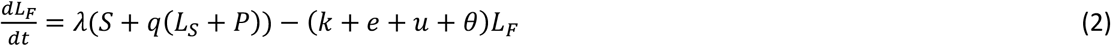

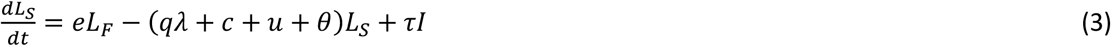

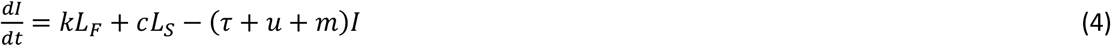

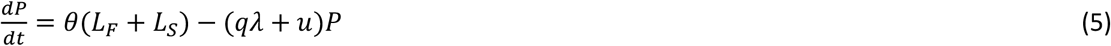

#### Model 2

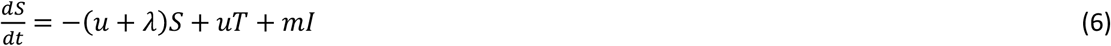

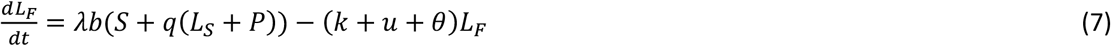

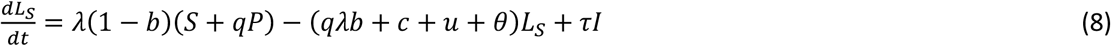

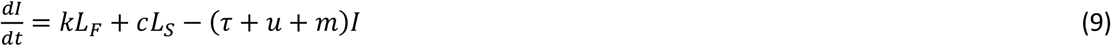

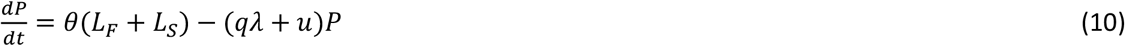

#### Model 3

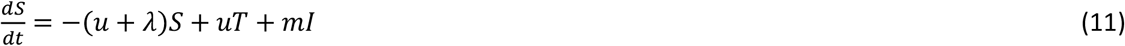

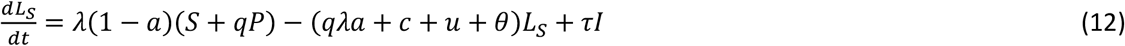

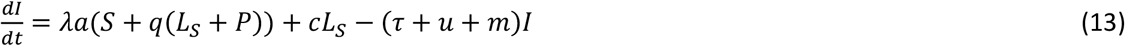

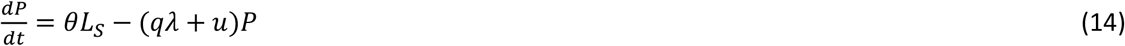

### Steady state solutions

For each model we can show that with reinfection (*q* > 0) but in the absence of preventive therapy (*θ* = 0) the steady state value of *I* is given by the solution to a quadratic equation of the form:

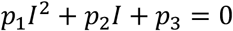

#### Model 1

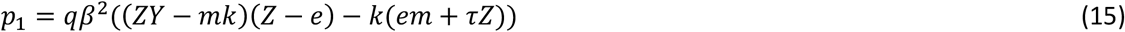

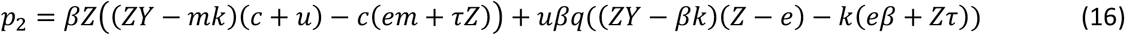

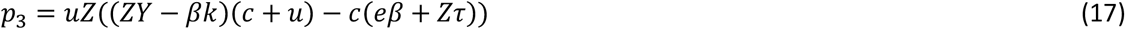

where

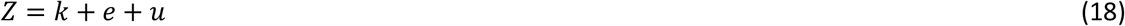

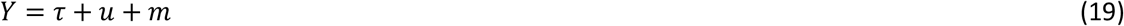

The other state variables are given by:

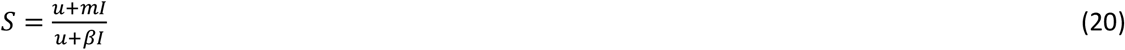

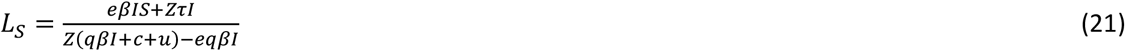

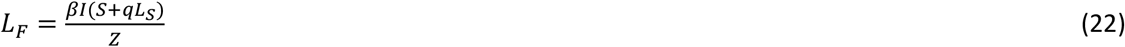

#### Model 2

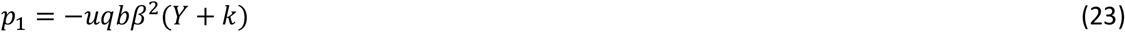

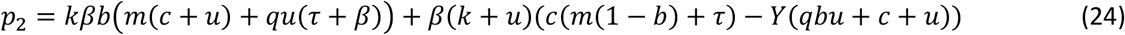

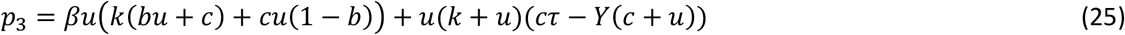

where

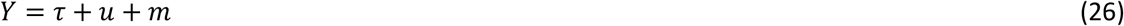

The other state variables are given by:

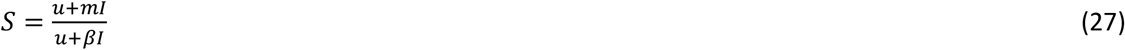

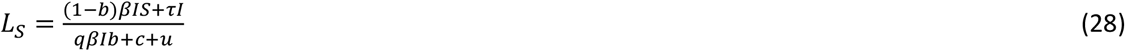

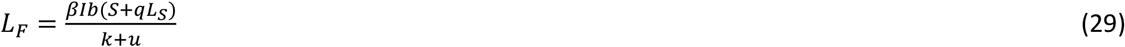

#### Model 3

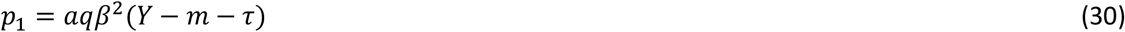

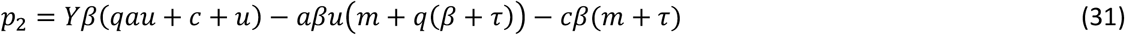

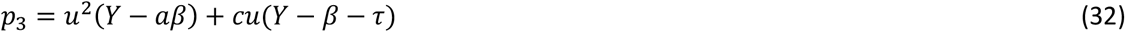

where

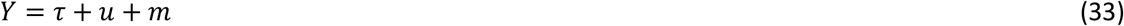

The other state variables are given by:

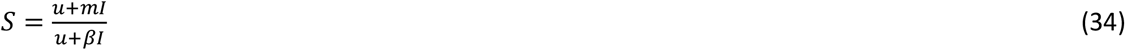

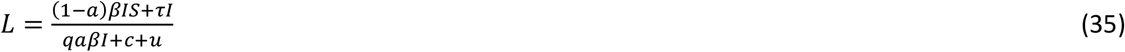

### Cumulative incidence of TB

The final proportion of infected individuals who will develop disease in model *i*, *P*_*i*_ is given by:

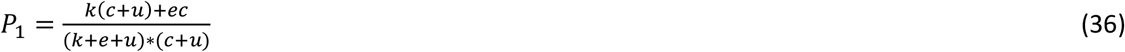

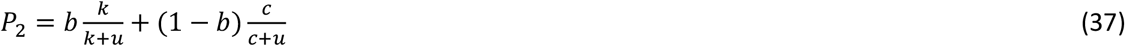

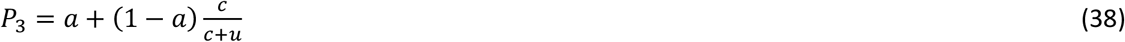

Rearranging equation (38) we can obtain an expression for the value of *a*, the proportion progressing directly to disease in model 3:

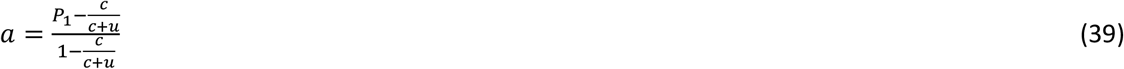

